# Designing a novel multi-epitope T vaccine for “targeting protein for Xklp-2” (TPX2) in hepatocellular carcinoma based on immunoinformatics approach

**DOI:** 10.1101/570952

**Authors:** Parisa Ghahremanifard, Farzaneh Afzali, Amin Rostami, Zahra Nayeri, Bijan Bambai, Zarrin Minuchehr

## Abstract

Hepatocellular carcinoma (HCC) is one of the leading cancer-related deaths worldwide. Recently, studies for HCC treatment are focused on cancer immunotherapy, particularly cancer vaccines, to complete and assist other therapies. TPX2 is a microtubule-associated protein necessary for cell division; therefore, alteration in its expression, especially up regulation, is associated with several human carcinomas such as HCC.

In this study, immunoinformatics tools were used to design a rational multi-epitope T vaccine against TPX2 in HCC. Cytotoxic T lymphocytes (CTL) and Helper T lymphocytes (HTL) epitopes were predicted and Maltose-binding protein (MBP) was added to the construct as an adjuvant. Evaluation of vaccine properties was indicated that our construct is stable and immunogenic enough to induce relevant responses besides not being allergic. After predicting the tertiary structure and energy minimization, protein-protein docking was performed to calculate the free energy of possible interactions between the vaccine and toll-like receptor 4 (TLR4) to assure that simultaneous complementary responses would be activated by our construct. Finally, Codon optimization and *in-silico* cloning were performed to ensure the vaccine expression efficiency in the desired host.

## Introduction

Among cancer-related death worldwide liver cancer is the second leading cause of 788,000 death in 2015 [1]. Hepatocellular carcinoma (HCC) is the most prevalent primary liver cancer which is commonly associated with cirrhosis. Factors lead to cirrhosis can be divided into two categories. Viral factors, including hepatitis B virus (HBV) and hepatitis C virus (HCV), and non-viral factors such as Alcohol and lifestyle (obesity) [2][3]. Traditional HCC treatments include surgery (i.e., liver resection and transplantation), radioactive seed implantation, trans-arterial chemoembolization (TACE) and radiofrequency ablation (RFA) do not lead to completely eliminate residual of cancer cells and even may lead to metastasis [4].

Current studies are focused on cancer immunotherapy, especially cancer vaccines, since they can activate cytotoxic T lymphocytes (CTL) therefore, it may be possible to eliminate cancerous cells without damaging the healthy ones [5].

Cell division needs a microtubule-associated protein known as *targeting protein for Xklp-2* (TPX2) [6]. Depletion of TPX2 can increase G2/M transition and rise cell polyploidy and genomic instability which in return induces multi-nucleation and DNA damage in HCC cells [7]. TPX2 expression is tightly controlled during cell cycle progression and its alterations have been reported to be associated with human carcinogenesis. TPX2 overexpression is reported in lung, cervical, bladder, esophageal, liver, and pancreatic cancer, [8]. Peptides derived from TPX2 can be recognized by CTLs, so in HCC replication cycle, TPX2 can be produced and recognized by CTLs, then HCC cells can be killed by activating CTLs [6] so it would be a rational candidate for designing anti-hepatocellular cancer vaccine.

Triggering Helper T lymphocytes (HTL) is a crucial step for the induction of CTL response [9]. Instead of using the whole protein as an antigen, only the immunogenic parts, i.e. epitopes, can be used for designing an immunogenic construct with ideal features [10]. In the present study, we used immunoinformatic tools to design a multi-epitope T vaccine including CTL and HTL epitopes to activate both cytotoxic and helper T lymphocytes. The ability of the vaccine construct in induction of the immune responses was also evaluated through multiple in silico assessments such as identifying its antigenicity and allergenicity, stability, and its capability in binding to immune receptors through protein-protein docking simulations.

## Methods

### 1. The followed steps in the current study

The main steps of designing the desired vaccine are shown in Fig. 1.

**Figure 1.**
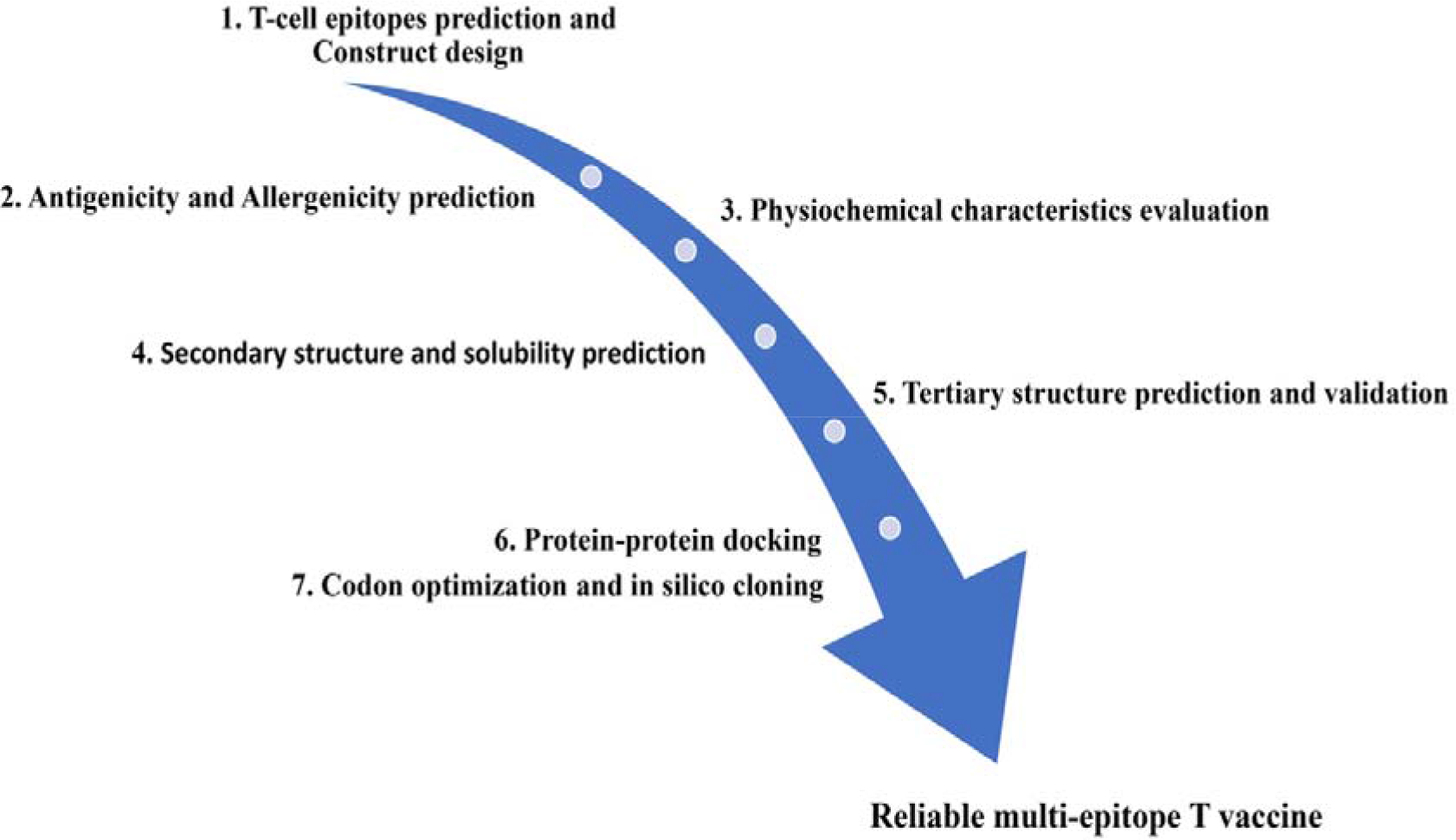
The summarized flowchart of designing multi-epitope T vaccine process.

### 2. Prediction of T-cell epitopes

Complete amino acid sequences of *Targeting protein for Xklp2* (UniProt ID: Q9ULW0) consists of 747 amino acids and Maltose/maltodextrin-binding periplasmic protein (UniProt ID: P0AEX9) consists of 396 amino acids were obtained from UniProt (http://www.expasy.org/uniprot).

In order to predict CTL epitopes for TPX2 several databases including Rankpep [11], IEDB [12], MHCPred [13] and Propred-I [14] were used. For evaluating the immunogenicity, these epitopes were submitted in IEDB MHC-I immunogenicity prediction module and the epitopes with highest scores that can induce CTL response were chosen. The HLA-A*02:01 was chosen for the super type [15]. For predicting HTL epitopes for TPX2, IEDB MHC-II epitope prediction module and NetMHCIIpan method were used. The HLA-DRB1*11:03 allele was chosen while the rest of the parameters were default [16]. The epitopes were sorted by their percentile rank. Lower percentile rank shows more affinity for binding to HTL receptor. Top 5 epitopes were selected with less percentile rank.

IFNepitope server (http://crdd.osdd.net/raghava/ifnepitope/predict.php) were used for evaluating the ability of selected HTL epitopes for inducing INF-γ production, IFN-γ versus other cytokine was chosen as a model of prediction while the rest of the parameters were as the default.

### 3. Designing the multi-epitope T construct and its improvement

CTL and HTL epitopes were fused by maximum immunogenic capability together. For inducing innate immune response as well as adaptive immune responses an adjuvant should be added to the construct. Maltose-binding protein (MBP) not only has intrinsic adjuvant-like properties [17] but also adding it at the N- or C-terminus of the protein can facilitate the purification process [18]. [19]. MBP was linked to N-terminal of the vaccine construct using EAAAK linker while HTL epitopes were linked together by the sequence of KFER. CTL epitopes were also linked together using the sequence of AK [20].

### 4. Antigenicity and Allergenicity prediction

ANTIGENpro (http://scratch.proteomics.ics.uci.edu/) and VaxiJen (0.4% threshold) (http://www.ddg-pharmfac.net/vaxijen/VaxiJen/VaxiJen.html) servers were used for antigenicity and allergenicity evaluation of final vaccine candidate. ANTIGENpro can classify proteins in the antigenic or non-antiganic group by using SVM and VaxiJen Prediction of antigen is alignment-free and based on various physiochemical properties of protein. AllerTOP v. 2.0 server (http://www.ddg-pharmfac.net/AllerTOP/) was used for the vaccine’s allergenicity prediction.

### 5. Evaluation of physicochemical characteristics of the vaccine

ProtParam server (https://web.expasy.org/protparam/) was used for evaluating physicochemical characteristic of the designed vaccine such as molecular weight, theoretical pI, instability index, aliphatic index, and grand average of hydropathicity (GRAVY). This server is an online tool which can calculate different physicochemical characteristic on the basis of pK values of different amino acids.

### 6. Secondary and Tertiary structure prediction, validation, and energy minimization

Secondary structure of the vaccine was predicted using GOR4 server (https://npsa-prabi.ibcp.fr/cgi-bin/npsa_automat.pl?page=/NPSA/npsa_gor4.html).

Tertiary structure of the vaccine construct was predicted using I-TASSER server (https://zhanglab.ccmb.med.umich.edu/I-TASSER/). I-TASSER works based on multiple-threading alignments and iterative template fragment assembly simulations to provide the most accurate structure [21]. For estimating the quality of predicted structure, the confident score (C-score) was used; higher C-score indicates higher confidence of the model. While TM-score and root mean square deviation (RMSD) are known standards for measuring the structural similarity between protein structures; lower values show high resolution and better fits of the model [22]. The ProSA-web online server (https://prosa.services.came.sbg.ac.at/prosa.php) was used to screen potential errors. The calculated z-score defines the accuracy of the model. Ramachandran plot of the RAMPAGE server (http://mordred.bioc.cam.ac.uk/~rapper/rampage.php) performed and obtained to define stereochemical quality of the final model.

### 7. Protein-protein docking of the vaccine with TLR4

The tertiary structures of vaccine construct and TLR4 (PDB ID: 3FXI) as a receptor of the attached adjuvant [23] was prepared by Chimera 1.11.2. protein-protein interactions were analyzed using HEX 8.0.0 docking software. The predicted tertiary structure of the vaccine candidate was experienced energy minimization by SPDBV 4.10 with Gromos96 force field before docking. The total free binding energy of interaction measured by regarding shape and electrostatics correlation type [24]. The solution was set to 1000, final search to 25, and other parameters were kept unchanged.

### 8. Codon optimization and *in-silico* cloning

For optimizing the Codon Usage of vaccine candidate for *E. coli*, K12 strain as the host, JCAT (http://www.jcat.de/) was used [25][26]. For *in-silico* cloning of the optimized vaccine candidate sequence in pET28a (+), XhoI and NdeI restriction sites were placed by using SnapGene tool which can simulate restriction cloning.

## Results

### 1. Selected CTL and HTL epitopes

Cytotoxic T lymphocytes are able to kill cancerous cells. For this purpose, we need immunogenic CTL epitopes which can induce CTL response [27]. In order to predict CTL epitopes of TPX2 several servers were used. 739 epitopes from Rankpep 739 from IEDB 515 from MHCPred and top ten epitopes from Propred-I with 9 mer length were obtained. The top 4 consensus epitopes among mentioned databases with higher IEDB MHC-I immunogenicity scores were selected for constructing the final vaccine candidate (Table. 1A). Top 10 CTL epitopes predicted by different servers and their immunogenicity score are listed in supplementary Table 1. For predicting HTL epitopes for TPX2, IEDB MHC-II epitope prediction tool was used. Top 4 epitopes which had less percentile rank and IC50 values with 15mer length were selected. (Table. 1B). Then for evaluating the ability to produce INF-γ for final epitopes IFNepitope server was used, and the result for all the epitopes was positive which shows they can activate Th1 and induce the production of INF-γ (Table. 1C). Top 10 predicted HTL epitopes and their score from the IFNepitope server are listed in supplementary Table 2.

**Table 1.**
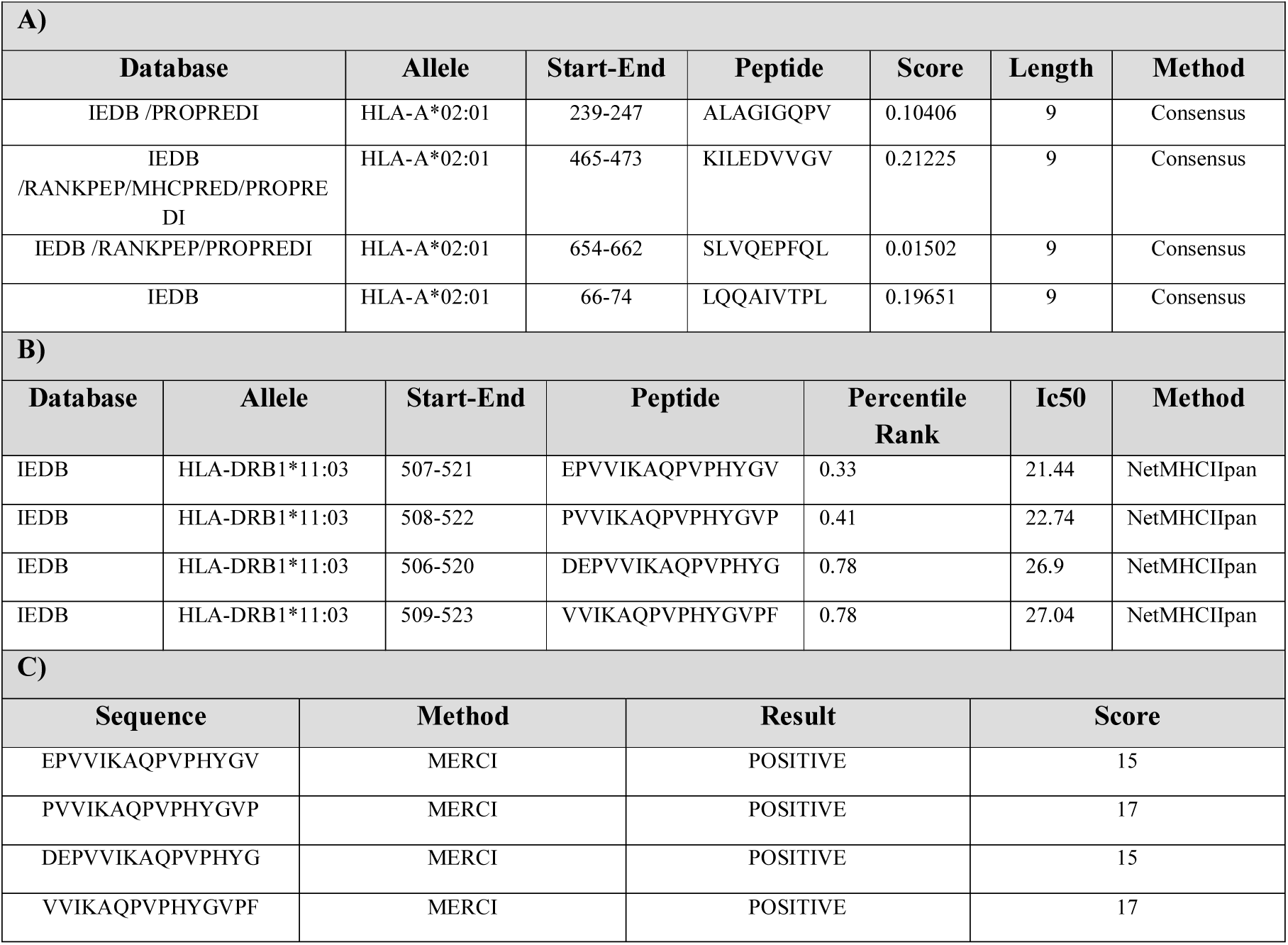
Selected CTL and HTL epitopes. **A)** Selected CTL and their immunogenicity score. **B)** Selected HTL epitopes and their percentile rank and IC50. **C)** Selected HTL epitopes and their score from the IFNepitope server.

### 2. Designing multi-epitope T construct

The final construct with 556 amino acids includes 5 CTL and 5 HTL epitopes accompanied by MBP sequence added to its N-terminal. All used linkers sequences are shown in Fig. 2A.

**Figure 2.**
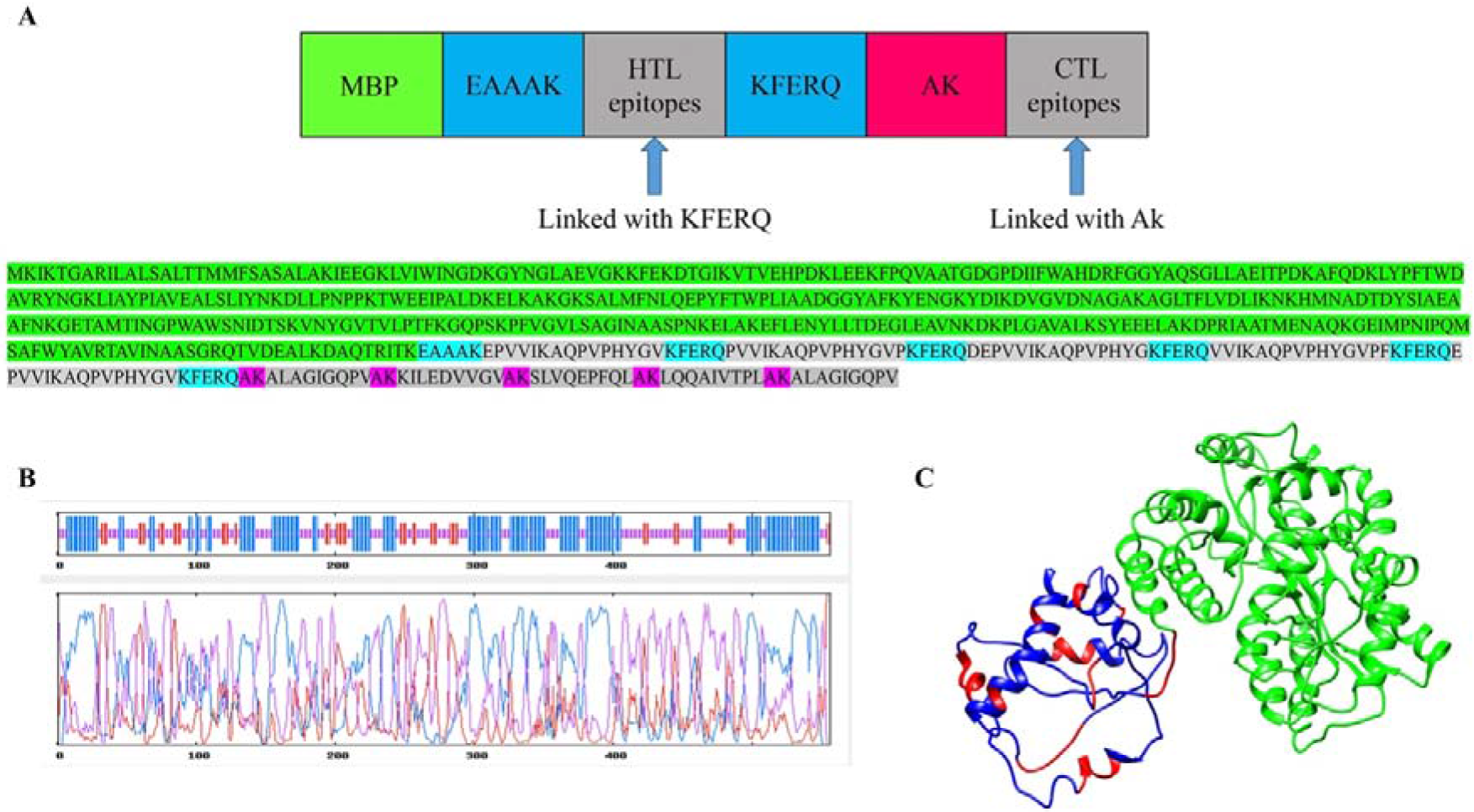
A) Schematic concept of the vaccine accompanied by sequence in below. The green sequence relates to MBP, cyan and pink relate to linkers, and epitopes are shown in grey. B) secondary structure of the vaccine helix (blue), sheet (red), and coil (purple). C) 3D structure of the vaccine: MBP (green), linkers (red), epitopes (blue).

### 3. Antigenicity and Allergenicity evaluation

Antigenicity of the vaccine predicted by ANTIGENpro and VaxiJen were respectively 0.80 and 0.43, which confirm the antigenicity of the vaccine construct [28][29].

Allergenicity of the vaccine was predicted by AllerTOP v. 2.0 server and the result defined it as non-allergen.

### 4. Physicochemical characteristics evaluation of the final vaccine

For evaluating physicochemical characteristics of the final vaccine ProtParam server was used. According to the data obtained from the server, the molecular weight of the vaccine candidate was 60.94 kDa; predicted pI was 8.62; and evaluated half-life was shown that the construct is stable Evaluated aliphatic index was 88.83 which indicates that the vaccine is thermostable. GRAVY was calculated as −0.219 demonstrating the hydrophilic nature of vaccine. The instability index score was 32.04, which classifies the vaccine as stable [30].

### 5. Secondary and Tertiary structure prediction, validation, and energy minimization

The secondary structure of the vaccine construct was predicted using GOR4 web tool. The result indicates that the secondary structure of the vaccine contains 43.88% random coil, 43.17% alpha helix, and 12.95% extended strands (Fig. 2B).

Among predicted five models of I-TASSER, first model with the highest C-score = − 1.2, TM-score = 0.56±0.15 and RMSD = 10.4±4.6Å was chosen as the best model (Fig. 2C).

The z-score obtained from ProSA-web before and after minimization by SPDBV 4.10 was −11.2 and −11.27 respectively (Fig. 3A) showing the reliability of the final construct in resembling native proteins. According to Ramachandran plot before minimization, 83.6% of residues were in the favored region, 12.8% in the allowed region and 3.6% in outlier region while the values were improved by 85.6%, 11.2%, and 3.2% respectively after minimization (Fig. 3B).

**Figure 3.**
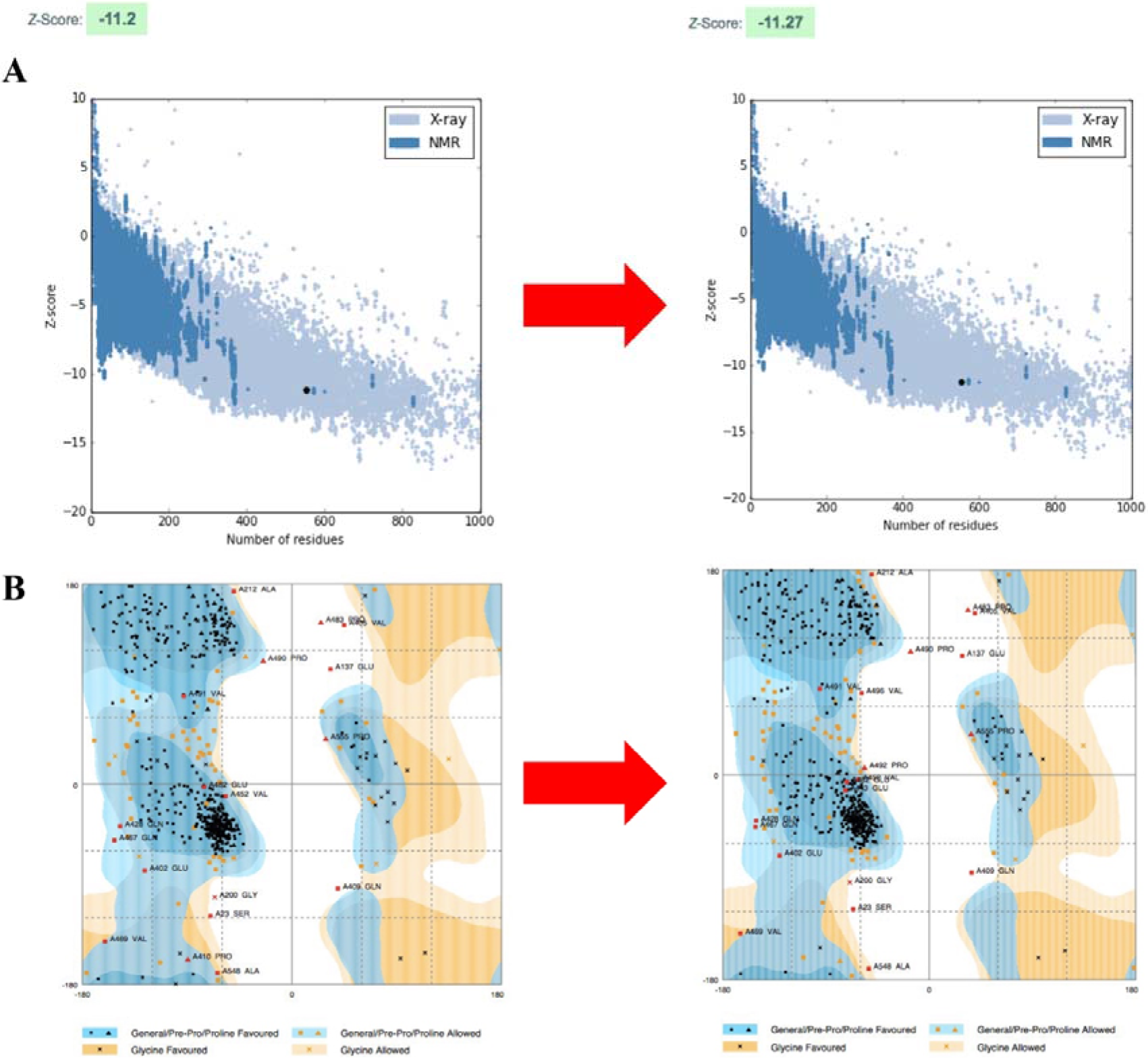
ProSA-web Z-score plot and Ramachandran plot before and after minimization. A) Z-score before and after minimization (the modelled construct shown as black spot) B) The improvement of the final construct through minimization visualized by Ramachandran plot.

### 6. Protein-protein docking of the vaccine with TLR4

Protein-protein docking was used for evaluating the interaction between vaccine candidate and TLR4. The estimated binding energy and pose of interaction are two crucial factors in this analysis. The pose of interaction is shown in (Fig. 4A) with −15601.8 as the resulted binding energy that shows the possibility of interaction between our designed vaccine and TLR4 which improves other complementary inflammatory responses against desired antigen.

**Figure 4.**
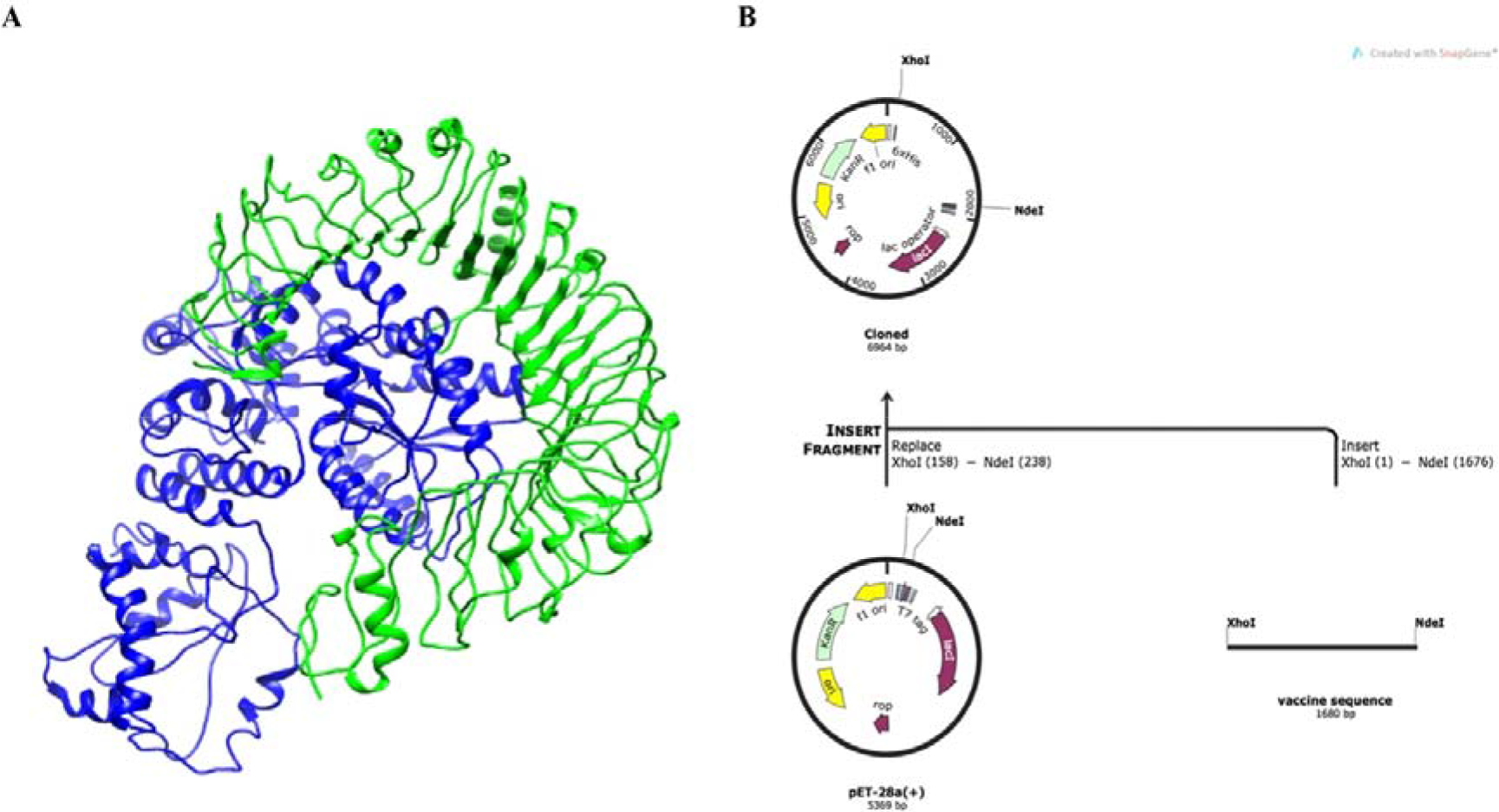
**A)** Docked complex of vaccine as a ligand in blue color and TLR4 as a receptor in green color. **B)** In silico cloning of the optimized vaccine sequence in to pET28a(+)

### 7. Codon optimization and in silico cloning

In order to reach the maximum level of expression in *Escherichia coli (E. coli)* k12 strain, the DNA sequence of the vaccine was optimized by Java Codon Adaptation Tool (JCAT). The CAI for the optimized vaccine sequence was 1 which is ideal, and GC content was 50 which is also in the perfect range (30%-70%) [28]. After adding restriction sites of XhoI and NdeI in the optimized sequence, it was cloned in pET28a(+)[31] using SnapGene tool (Fig. 4B) [28][9]. The Total length of the vector containing our construct is 6964 bp.

## Discussion

Worldwide, about 10% of all deaths - in adults - is associated with HCC. Unfortunately, it is diagnosed too late, and also may lead to metastasis because of the lack of proper treating procedures [32]. Hence, new therapies for solving this severe problem are emerging. Recently, studies for HCC treatment are focused on cancer immunotherapy especially cancer vaccines. Unlike traditional vaccines which prevent a certain disease, cancer vaccines are designed to treat existing disease [33]. Among various types of vaccines, synthetic peptide vaccines are highly specific and easy to produce [34].

Whether a vaccine is successful or not highly depends on the tumor antigen targeted by the vaccine. The overexpressed self-antigens, as the name indicates, are overexpressed in tumors, while their presence is low but detectable in normal cells. These antigens induce immune responses only against cancerous cells and leave normal ones unharmed, in other words, they do not provoke autoimmunity [35] [36].

In this study we have designed an anti-HCC peptide vaccine based on TPX2 that is included in overexpressed self-antigens group and is overexpressed in several human carcinomas. With reference to Chao-Wen Hsu et al, TPX2 is overexpressed in HCC, hence counts as a potential target for immunotherapy against it [7] [8].

An appropriate synthetic peptide cancer vaccine should be able to induce CTL responses in order to kill the cancerous cells, which require cytokines that are produced by TH cells [37]. Therefore, it needs to induce both CTL and HTL responses, according to that we tried to predict CTL and HTL epitopes using several immunoinformatic databases. Nowadays immunoinformatic can play a crucial role in designing a cancer vaccine by providing efficient methods for finding a potential candidate for immunotherapy by compering epitopes to find the best candidate based on MHC-binding affinity of the epitopes which are capable to induce a strong immune response[38]. Reports indicate that experimental vaccine discovery and development takes approximately 10-20 years and cost of hundreds of million US dollars. While with the help of immunoinformatic tools we can save in time and cost [39]. After predicting CTL and HTL epitopes the ones with less percentile rank which show higher binding affinity were selected. Also, the immunogenicity of the epitopes was evaluated and the result was satisfying.

Cancerous cells may suppress the immune response by producing factors such as (IL-10, TGF, VEGF, and etc.) thus escaping from immune system. To surmount the problem, adjuvants would be added to vaccine construct for generating CTLs which are capable of overcoming the suppressive mechanisms [40]. So, we considered it and applied it to our design and MBP sequence was added to N-terminal as an adjuvant to avoid the immune escape mechanisms. We chose MBP as an adjuvant not only because it is able to increase the immunogenicity of the construct but also it can facilitate the expression and purification process of recombinant protein in *E. coli*, by one-step purification using affinity chromatography (cross-linked amylose) [18][23].

The results for antigenicity and allergenicity of the vaccine construct showed that this vaccine is able to induce a strong immune response while it was known as non-allergen and safe to use. Physicochemical characteristics of the vaccine showed that it is hydrophilic and its structure is stable. The result for thermostability was also satisfying.

The Validity of the interaction between vaccine and receptor (TLR4) was evaluated by molecular docking. Finally, codon optimization and *in silico* cloning was performed to ensure the efficiency of expression of the vaccine in appropriate host.

However, for future use of this vaccine *in vivo* study need to be done for evaluating the immune response against HCC in the real condition. All in all, immunotherapies like monoclonal antibodies, CAR T-cell therapy, checkpoint inhibitors and cancer vaccines are less toxic than chemotherapeutic agent but they may have some side effects so they need further evaluation [41].

## Abbreviations

HCC: Hepatocellular carcinoma
MBP: Maltose-binding protein
HBV: hepatitis B virus
HCV: hepatitis C virus
TACE: transarterial chemoembolization
RFA: radiofrequency ablation
CTL: cytotoxic T lymphocytes
TPX2: targeting protein for Xklp-2
HTL: helper T lymphocytes
TSA: tumor-specific antigens
TAA: tumor-associated antigens
MHC: major histocompatibility complex
GRAVY: Grand average of hydropathicity
E. coli: Escherichia coli
JCAT: Java Codon Adaptation Tool
CAI: Codon Adaptation Index
RMSD: root mean square deviation

## Author Contributions statement

Z.M, B.B and P.GH conceptualized the project. F.A and P.GH did the analyses and interpreted the data except the data related to docking part which was done by Z.N and F.A. P.GH and A.R wrote the manuscript. Z.M supported the project (corresponding author). All authors read and approved the final manuscript.

## Acknowledgments

The financial support of this research project by the National Institute of Genetic Engineering and Biotechnology (NIGEB) of Iran is acknowledged.

