## supplementary Table 1-supplementary Table 2- for "Designing a novel multi-epitope T vaccine for “targeting protein for Xklp-2” (TPX2) in hepatocellular carcinoma based on immunoinformatics approach"

**Supplementary Table 1:** Top 10 predicted CTL epitopes by different servers and their immunogenicity score. A) RANKPEP - B) IEDB MHC-I Binding - C) MHCPRED - D) PROPREDI

**A)** **RANKPEP**

| **RANK** | **POS.** | **N** | **SEQUENCE** | **C** | **MW (Da)** | **SCORE** | **% OPT.** | **Class I Immunogenicity** |
| --- | --- | --- | --- | --- | --- | --- | --- | --- |
| 1 | 250 | VKK | SVSQVTKSV | DFH | 916.03 | 85 | 66.41% | -0.37218 |
| 2 | 673 | ERQ | ELEKRMAEV | EAQ | 1086.28 | 84 | 65.62% | -0.20788 |
| 3 | 465 | CPT | KILEDVVGV | PEK | 953.14 | 82 | 64.06% | 0.21225 |
| 4 | 352 | DDI | NLLPSKSSV | TKI | 926.08 | 80 | 62.50% | -0.61514 |
| 5 | 154 | IDE | ILPSKKMKV | SNN | 1025.35 | 79 | 61.72% | -0.85727 |
| 6 | 654 | LSG | SLVQEPFQL | ATE | 1042.21 | 77 | 60.16% | 0.01502 |
| 7 | 220 | KSM | KMQQEVVEM | RKK | 1103.31 | 74 | 57.81% | 0.07554 |
| 8 | 69 | LQQ | AIVTPLKPV | DNT | 919.17 | 74 | 57.81% | -0.15726 |
| 9 | 246 | IGQ | PVKKSVSQV | TKS | 953.14 | 71 | 55.47% | -0.61654 |
| 10 | 296 | SPA | RVTKGCTIV | KPF | 958.17 | 65 | 50.78% | -0.11163 |

**B)** **IEDB MHC-I Binding**

| **RANK** | **start** | **end** | **peptide** | **method** | **percentile rank** | **ann_ic50** | **ann_rank** | **Class I Immunogenicity** |
| --- | --- | --- | --- | --- | --- | --- | --- | --- |
| 1 | 239 | 247 | ALAGIGQPV | Consensus | 0.4 | 16.01 | 0.15 | 0.10406 |
| 2 | 465 | 473 | KILEDVVGV | Consensus | 0.5 | 7.67 | 0.06 | 0.21225 |
| 3 | 352 | 360 | NLLPSKSSV | Consensus | 2 | 259.46 | 2 | -0.61514 |
| 4 | 53 | 61 | GLFQGKTPL | Consensus | 2.3 | 48.87 | 0.53 | -0.22228 |
| 5 | 631 | 639 | TVISQEPFV | Consensus | 2.3 | 319.6 | 2.3 | -0.08278 |
| 6 | 654 | 662 | SLVQEPFQL | Consensus | 2.3 | 78.84 | 0.79 | 0.01502 |
| 7 | 396 | 404 | KLQQYKFKA | Consensus | 2.6 | 308.23 | 2.2 | -0.38796 |
| 8 | 66 | 74 | LQQAIVTPL | Consensus | 3.5 | 319.43 | 2.3 | 0.19651 |
| 9 | 154 | 162 | ILPSKKMKV | Consensus | 3.5 | 442.3 | 2.8 | -0.85727 |
| 10 | 410 | 418 | RILEGGPIL | Consensus | 4.4 | 370.5 | 2.5 | 0.23045 |

**C) MHCPRED**

| **RANK** | **peptide** | **Predicted -logIC50 (M)** | **Predicted IC50 Value (nM)** | **Confidence of prediction (Max = 1)** | **Class I Immunogenicity** |
| --- | --- | --- | --- | --- | --- |
| 1 | [KILEDVVGV](http://www.ddg-pharmfac.net/mhcpred/scripts/MHCPred_scripts/additive.pl) | 7.539 | 28.91 | 1 | 0.21225 |
| 2 | [DDINLLPSK](http://www.ddg-pharmfac.net/mhcpred/scripts/MHCPred_scripts/additive.pl) | 7.485 | 32.73 | 0.78 | -0.09057 |
| 3 | [PKFKALPLP](http://www.ddg-pharmfac.net/mhcpred/scripts/MHCPred_scripts/additive.pl) | 7.475 | 33.5 | 0.78 | -0.16718 |
| 4 | [KKRTFDETV](http://www.ddg-pharmfac.net/mhcpred/scripts/MHCPred_scripts/additive.pl) | 7.471 | 33.81 | 0.89 | 0.29792 |
| 5 | [PILPKKPPV](http://www.ddg-pharmfac.net/mhcpred/scripts/MHCPred_scripts/additive.pl) | 7.466 | 34.2 | 1 | -0.4436 |
| 6 | [FLKSTEEQE](http://www.ddg-pharmfac.net/mhcpred/scripts/MHCPred_scripts/additive.pl) | 7.466 | 34.2 | 0.89 | -0.0876 |
| 7 | [IIDEILPSK](http://www.ddg-pharmfac.net/mhcpred/scripts/MHCPred_scripts/additive.pl) | 7.46 | 34.67 | 0.89 | 0.12109 |
| 8 | [INFRKLPSH](http://www.ddg-pharmfac.net/mhcpred/scripts/MHCPred_scripts/additive.pl) | 7.456 | 34.99 | 0.78 | -0.23638 |
| 9 | [IVKPFNLSQ](http://www.ddg-pharmfac.net/mhcpred/scripts/MHCPred_scripts/additive.pl) | 7.397 | 40.09 | 0.89 | -0.07927 |
| 10 | [KALPLPHFD](http://www.ddg-pharmfac.net/mhcpred/scripts/MHCPred_scripts/additive.pl) | 7.379 | 41.78 | 0.89 | 0.0597 |

**D) PROPREDI**

| **RANK** | **peptide** | **Position** | **Real Score** | **Log Score** | **% of Highest on log scale** | **Class I Immunogenicity** |
| --- | --- | --- | --- | --- | --- | --- |
| 1 | KILEDVVGV | 465 | 1167.868063 | 7.0629 | 39.55 | 0.21225 |
| 2 | NLLPSKSSV | 352 | 257.3424 | 5.5504 | 31.08 | -0.61514 |
| 3 | SLVQEPFQL | 654 | 123.90192 | 4.8195 | 26.99 | 0.01502 |
| 4 | ILPSKKMKV | 154 | 118.2384 | 4.7727 | 26.73 | -0.85727 |
| 5 | KLQQYKFKA | 396 | 100.8504 | 4.6136 | 25.84 | -0.38796 |
| 6 | GLFQGKTPL | 53 | 79.04088 | 4.37 | 24.47 | -0.22228 |
| 7 | ALAGIGQPV | 239 | 69.552 | 4.2421 | 23.76 | 0.10406 |
| 8 | ALPLPHFDT | 568 | 43.2216 | 3.7663 | 21.09 | 0.11665 |
| 9 | TVISQEPFV | 631 | 33.4719 | 3.5107 | 19.66 | -0.08278 |
| 10 | KMQQEVVEM | 220 | 28.8834 | 3.3633 | 18.84 | 0.07554 |

Class I Immunogenicity positive

Selected epitopes

**Supplementary Table 2:** Top 10 predicted HTL epitopes and their score from the IFNepitope server

**B)** **IEDB MHC-I Binding**

| **RANK** | **start** | **end** | **peptide** | **method** | **percentile rank** | **netmhciipan_ic50** | **netmhciipan_rank** | **IFNepitope server** |
| --- | --- | --- | --- | --- | --- | --- | --- | --- |
| 1 | 507 | 521 | EPVVIKAQPVPHYGV | NetMHCIIpan | 0.33 | 21.44 | 0.33 | POSITIVE |
| 2 | 508 | 522 | PVVIKAQPVPHYGVP | NetMHCIIpan | 0.41 | 22.74 | 0.41 | POSITIVE |
| 3 | 506 | 520 | DEPVVIKAQPVPHYG | NetMHCIIpan | 0.78 | 26.9 | 0.78 | POSITIVE |
| 4 | 509 | 523 | VVIKAQPVPHYGVPF | NetMHCIIpan | 0.78 | 27.04 | 0.78 | POSITIVE |
| 5 | 505 | 519 | EDEPVVIKAQPVPHY | NetMHCIIpan | 1.09 | 31.97 | 1.09 | POSITIVE |
| 6 | 707 | 721 | LRRELVHKANPIRKY | NetMHCIIpan | 1.45 | 35.84 | 1.45 | POSITIVE |
| 7 | 708 | 722 | RRELVHKANPIRKYQ | NetMHCIIpan | 1.69 | 38.41 | 1.69 | POSITIVE |
| 8 | 706 | 720 | RLRRELVHKANPIRK | NetMHCIIpan | 2.53 | 46.06 | 2.53 | POSITIVE |
| 9 | 709 | 723 | RELVHKANPIRKYQG | NetMHCIIpan | 2.68 | 48.2 | 2.68 | POSITIVE |
| 10 | 510 | 524 | VIKAQPVPHYGVPFK | NetMHCIIpan | 3.23 | 54.07 | 3.23 | POSITIVE |

Selected epitopes
